# Stimulation-mediated reverse engineering of silent neural networks

**DOI:** 10.1101/2021.04.14.439683

**Authors:** Xiaoxuan Ren, Aviad Hai

## Abstract

Reconstructing connectivity of neuronal networks from single cell activity is essential to understanding brain function, but the challenge of deciphering connections from populations of silent neurons has been largely unmet. We demonstrate a protocol for deriving connectivity of realistic silent neuronal networks using stimulation combined with a supervised learning algorithm, that enables inferring connection weights with high fidelity and predicting spike trains at the single-spike and single-cell level with high accuracy. These testable predictions about the number and protocol of the required stimulations is expected to enhance future efforts for deriving neuronal connectivity and drive new experiments to better understand brain function.

## 1. Introduction

Deciphering the function of neurobiological networks hinges on proper determination of connectivity at the single neuronal level[1–4]. Uncovering the specific circuit connectivity and how it mediates neuronal firing can then lead to more effective therapies for brain disorders[5–10]. Different methodologies have been used to derive neural coupling at the neuronal level[11–16] and at the brain regional level [17–19]. Current efforts involve reconstructing highly-detailed structural connections [11, 20, 21], or deriving putative mono-synaptic connections from large-scale electrophysiological recordings[15, 22] and optical imaging [2, 16, 23, 24]. Notwithstanding the growing availability and increased precision of neurological imaging modalities, multiplexed electrophysiological recordings of action potentials by microelectrode arrays (MEAs) are still the mainstay input for decoding connectivity at the microcircuit resolution[14, 25, 26]. MEAs provide readouts at increasingly larger scales, with recent probes able to provide *in vivo* recordings of up to thousands of units in parallel[27–30].

Most studies rely primarily on active units to derive connectivity, and largely exclude non-firing or minimally active neurons. However, numerous groups present evidence that more than half of the neurons in the brain are “silent”, firing at very low frequencies [31, 32] and having highly specialized receptive fields [33–35]. Minimally active neurons do not fire spontaneously or participate in oscillatory activity, but instead usually fire at rates of 1 spike/min or less [36–38], with some groups reporting on frequencies of as low as 0.001 Hz [39] depending on the brain region and task studied. The proportion of silent neurons in a typical brain tissue varies, ranging between at least 10-20% in the sensory cortex[38, 39] to as much as 66% in the motor cortex [36]. Low firing rates limit the accuracy of network reconstruction, due to the inherently low information content at baseline conditions. Electrical stimulation[40, 41], sensory stimulation[42, 43] and neuropharmacological manipulations[44–46] are often used in neurobiological research to perturb a system above baseline, and could therefore activate silent neurons and thus improved the inference of network connectivity.

In this work we demonstrate a supervised learning method to analyze the use of stimulation for determining connectivity between postsynaptic neurons and heterogenous populations of both silent and active presynaptic neurons. We show that repetitive stimulation epochs can evoke activity that is sufficient for determining the synaptic weights of the entire neuronal population. We then characterize the performance of our algorithm for determining weights from excitatory, inhibitory and unconnected neurons in the population, and compare the ability of the method to predict spike trains. This approach presents a new platform to increase the performance of learning algorithms for reconstructing neuronal networks, and for using neuronal stimulation to decipher large-scale brain circuitry.

## 2. Methods

### 2.1 Simulated neuronal population spike trains

We generated spike trains for populations of neurons (size ranging between 200 and 2000 cells) recorded during stimulus response. 66% of the spike trains corresponded to “silent” neurons (firing at 1 spike/min, or 0.017 Hz, [35, 47]) and the rest corresponded to responsive neurons (firing at 20 Hz [48, 49]). Spikes were generated by sampling randomly using a uniform distribution, with a probability of 1/f. In addition to the above realistic condition, we compared results using simulated spike trains from a population with no silent neurons, where all the 200 units fired at 20 Hz (hyperactive non-realistic condition).

### 2.2 Postsynaptic model neuron and connectivity

We simulated a postsynaptic integrate-and-fire neuron receiving inputs from 100 out of the total population spike trains. 80 inputs were set to be excitatory (having positive weights), and 20 inputs were set to be inhibitory (having negative weights) [50]. Input weights ranged between [−8, 8] mV, sampled randomly using a uniform distribution. Spikes in the integrate-and-fire neuron model were generated by summation of the inputs at each time step (1 ms intervals) using equation 1:

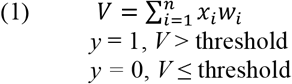

*V* is the membrane potential of the postsynaptic neuron, *x_i_* is the input from population neuron i, *w_i_* is the weight of input i, n = total number of neurons, threshold = 20 mV (with resting membrane potential = 0 mV), *y* is the spike output of the postsynaptic neuron.

### 2.3 Deriving connectivity

We used the simulated spike trains of the postsynaptic neuron and the population as ground-truth data for deriving connection weights using a perceptron algorithm[51, 52]. At each iteration of the algorithm, the derived connection weights were updated using the perceptron learning rule:

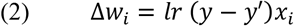

Δ*w_i_* is the update for input weight i, *lr* is the learning rate, *y* is the spike output of the postsynaptic neuron in the training data, and *y*′ is the output guessed by the perceptron model. We used 2500 trials (each of 1000 ms duration) to train the perceptron model. We used a learning rate of 0.01.

### 2.4 Performance measures

We evaluated the performance of the algorithm for deriving connectivity by the root mean squares error (RMSE) between the derived weights and actual input weights. In addition, we evaluated the performance by using the derived weights to predict spikes, as measured by the average of the prediction sensitivity and precision. Sensitivity was the ratio of true positives and actual spikes. Precision was the ratio of true positives and total predicted spikes. We tested the algorithm performance using 10 randomized simulations, with different spike trains and connectivity. The test data consisted of 2500 trials of 1000 ms each.

### 2.5 Stimulated network condition

We examined the effect of stimulation on deriving connectivity, by setting 50 of the population spike trains (cells chosen at random) to fire at 60 Hz during each stimulation duration (200 ms) based on commonly used parameters [53]. We examined results as a function of the number of stimulations, ranging from 1 to 15.

### 2.6 Connection types

We classified different connection types based on their weight, according to table 1. We classified weights as unconnected using a small ε (0.16 mV), which was the RMSE of derived weights for actual unconnected neurons in perceptron models that had high accuracy (performance > 0.99). We calculated the accuracy of classification for each connection type as the probability of synaptic weight *w* of being correctly classified, given the actual weight.

## 3. Results

### 3.1 prediction of synaptic weights for populations with silent neurons is improved by stimulation

We characterized the effect of silent neurons on deriving synaptic weights from a heterogenous population (excitatory, inhibitory and unconnected) to a postsynaptic neuron within an arena of a recording electrode (Fig. 1a,b), and characterized the improvement afforded by stimulation (Fig. 1c). As a reference case, we used a hyperactive neuronal population as a non-realistic scenario without silent neurons, for which the algorithm derived synaptic weights with high accuracy (Fig. 1d, RMSE = 0.01 ± 0.00, Pearson correlation r = 1, *p* = 0, n = 10 random populations). For the realistic neuron population (with silent neurons), the algorithm fails to predict the weights from silent neurons, and also from many of the active neurons (Fig. 1e, RMSE = 3.59 ± 0.42, Pearson correlation r = 0.44, *p* < 0.01). When the population was sufficiently stimulated, the algorithm was able to derive weights with high accuracy (Fig. 1f, RMSE is 0.35 ± 0.06, Pearson correlation r = 0.97, *p* ~ 0, 15 stimulations, 100 iterations, see supplementary Fig. 1). The stimulation protocol sufficiently improved the weight prediction of a population with silent neurons (Fig. 1g).

**Figure 1.**
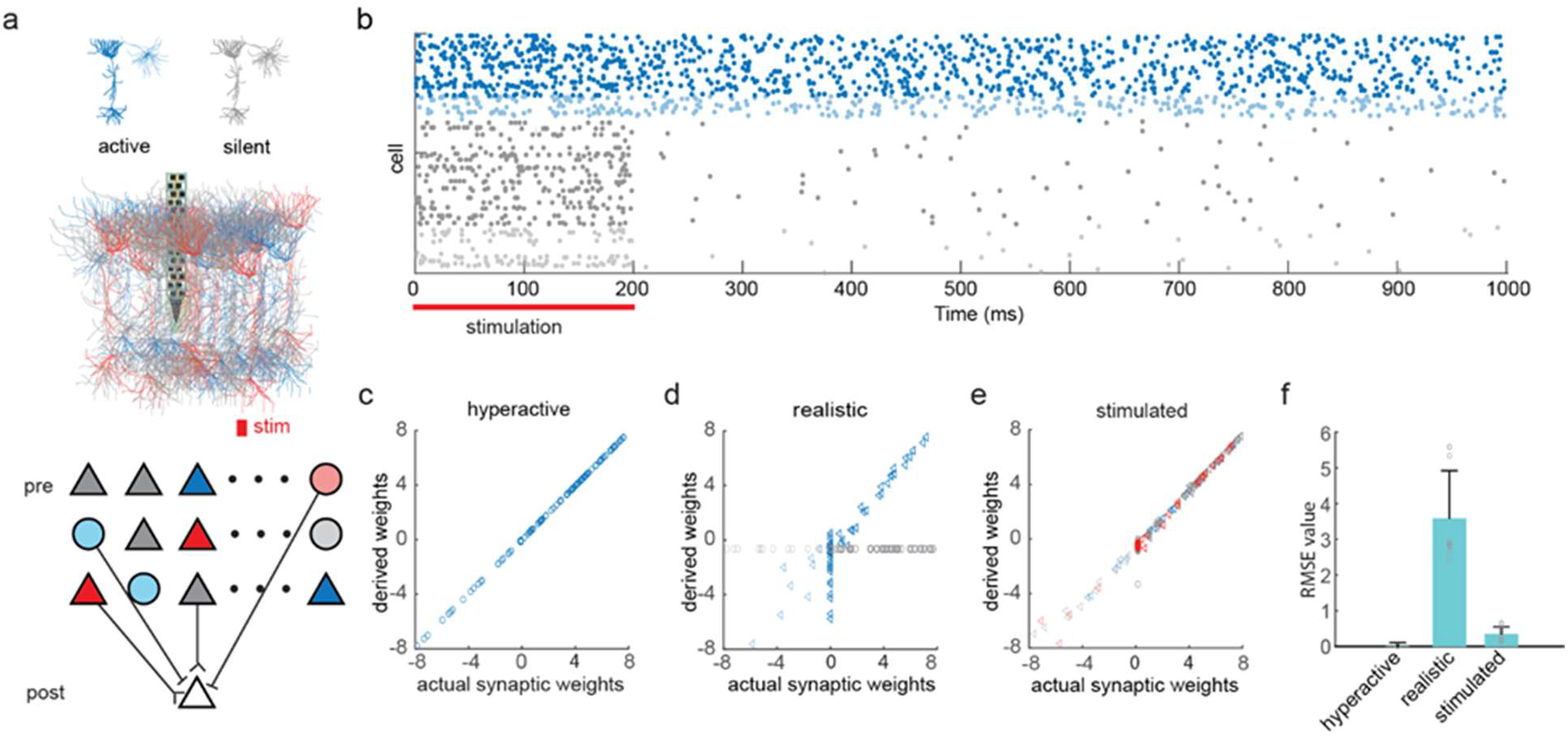
Stimulation-mediated derivation of synaptic weights from a neuronal population with silent cells. **(a)** Schematic diagram of the neuronal population (200 neurons, represented as spike trains) and its connections with an integrate-and-fire postsynaptic neuron. 66% of the neurons in the population were silent (grey, firing rate < 0.01 Hz) and the rest were active (blue, firing rate ~20 Hz). In the stimulated condition, 25% of the neurons were stimulated (red, firing rate ~60 Hz). The color-coding is maintained for the rest of the figure. The postsynaptic neuron received 80 excitatory inputs and 20 inhibitory inputs. **(b)** Spike raster of the population spike trains over 1000 ms. The top 34% of the trains correspond to active neurons firing at a baseline of 20Hz. The bottom 66% correspond to silent neurons that fire 1 spike/min. Stimulation duration was 200 ms. **(c)** Derived vs. actual weights in the hyperactive neuron population case (with no silent neurons). The derived weights matched the actual weights (Pearson correlation, r = 1.00, *p* ~ 0). **(d)** Derived vs. actual weights in the realistic neuron population case, where 66% of the population are silent. The derived weights deviated considerably from the actual weights (r = 0.44, *p* < 0.01). **(e)** Derived vs. actual weights in the stimulated population case, where 25% of the population were stimulated to fire at 60 Hz for 100ms. The derived weights closely matched the actual weights (r = 0.97, *p* ~ 0). **(f)** RMSE of the derived weights vs. actual weights for the cases shown in d-f (n=10), error bar is standard error of the mean.

**Figure 2.**
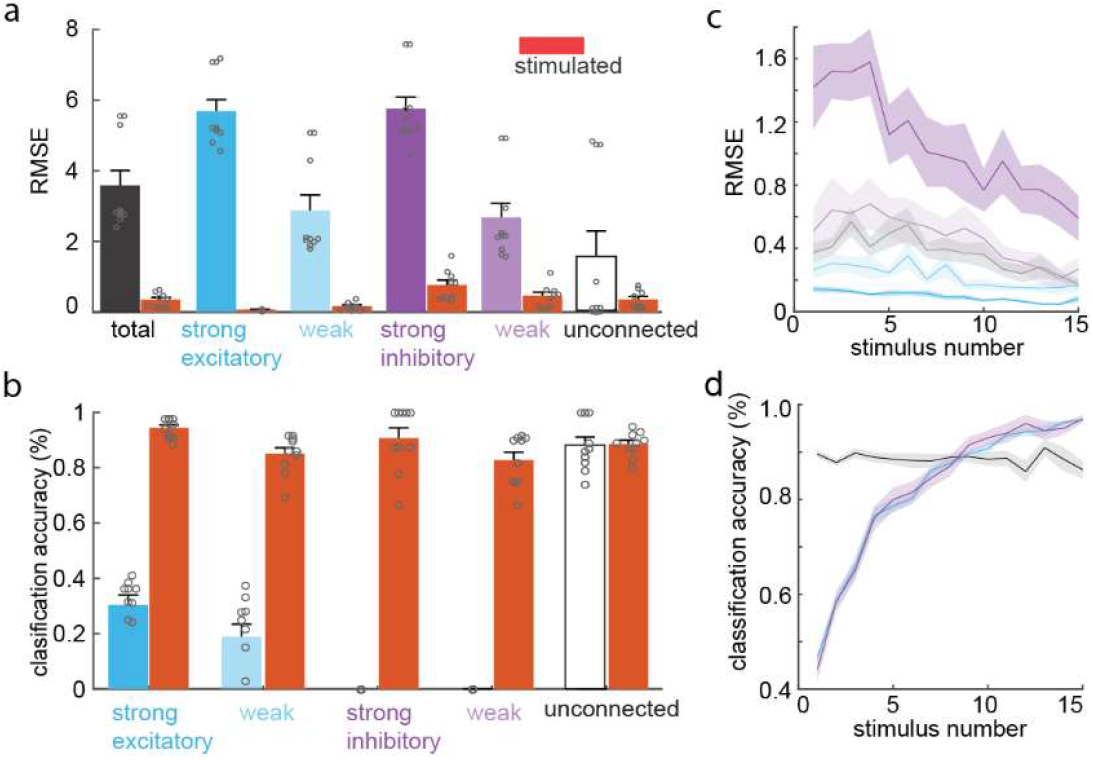
Deriving synaptic weights as a function of stimulus and connection type. **(a)** Performance of deriving connections of different type (excitatory and inhibitory) and strength (see Methods), in the unstimulated and stimulated (red) conditions. Stimulation improved performance for all cases except the unconnected (paired t-test, *p* < 0.05). **(b)** Classification accuracy for each connection type in the unstimulated and stimulated conditions. **(c)** Performance of deriving connection weights as a function of stimulus number, for the different connection types as shown in (a). The shaded area around each curve shows the standard deviation. **(d)** Classification accuracy for excitatory, inhibitory and unconnected neurons as a function of stimulus number.

### 3.2 Spike prediction using the derived weights

We validated the performance of the derived connection weights from unstimulated and stimulated populations in predicting spikes of the postsynaptic neuron, using test data of the population spikes in the stimulated condition (Fig. 3a). Weights derived from non-stimulated populations predicted spikes poorly, with a high proportion of false negatives and some false positives. In contrast, derived weights from stimulated populations predicted postsynaptic spikes with high accuracy, with the sensitivity and performance of the model reaching 0.96 ± 0.01 (Fig. 3b) and 0.97 ± 0.01 (Fig. 3c) after 15 stimuli, respectively. We also verified that the derived weights performed well on population data from the fully activated condition (Supplementary Fig. 2)

**Figure 3.**
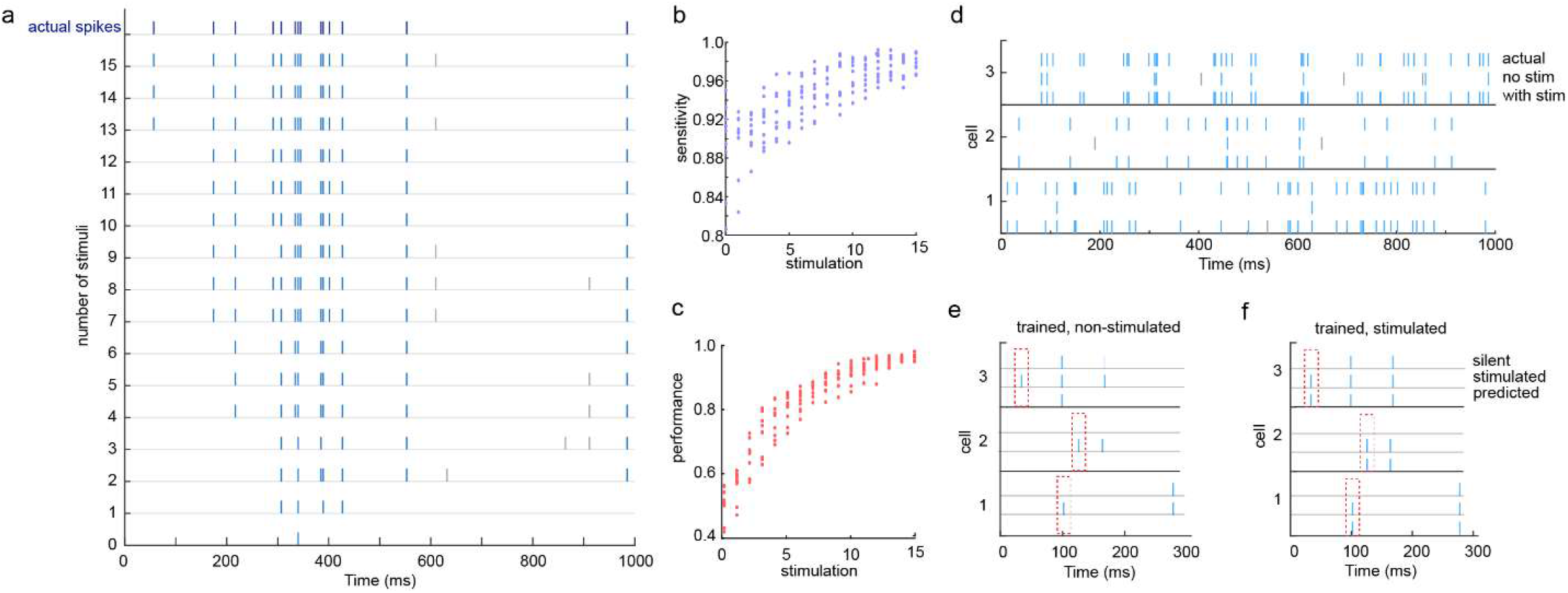
Spike prediction using the derived connection weights. **(a)** Predicted spikes with weights derived using different number of stimuli. The top spike train shows the actual postsynaptic spikes. Listed below, are spike trains predicted from populations stimulated with 0 to 15 stimuli. True positives are shown in blue, false positives are shown in grey. **(b)** Sensitivity of spike prediction as a function of stimulus number. The shaded area around the curve shows the standard deviation (n = 10). **(c)** Precision of spike prediction as a function of stimulus number. **(d)** Three examples of spike trains of the postsynaptic neuron in the test data (top), predicted spikes using derived weights from the non-stimulated population (middle) and stimulated population (bottom). Blue spikes are true positives, grey spikes are false negatives. **(e)** Three examples of spike trains of the postsynaptic neuron in the non-stimulated case (top), when stimulating one strong excitatory silent neuron (middle), and the predicted spikes using weights derived from the non-stimulated population data (bottom). **(f)** same as (e) but with predicted spikes using weights derived from the stimulated population data.

To verify that the stimulation of silent neurons significantly affects the postsynaptic spike pattern, we stimulated a single strong excitatory silent neuron, and compared the predicted postsynaptic spikes using weights derived from non-stimulated and stimulated data. Activating the single silent neuron was sufficient to trigger additional spikes (Fig. 3e), which were not predicted when using weights derived from non-stimulated population data (Fig. 3e), but were predicted faithfully using weights derived from stimulated population data (Fig. 3f).

To evaluate the accuracy of inferred weights of different connection types in realistic networks before and after stimulation, we compared the RMSE value of five groups of connections defined according to their type (excitatory or inhibitory) and strength (weak, strong and unconnected). The RMSE was improved by stimulation for all types of synaptic weights (Fig 2a, paired t-test: *p* < 0.05), except for the unconnected since their error was small to begin with (*p* = 0.1). Similarly, the classification accuracy of connection types (see Methods) improved by stimulation for all types, especially for inhibitory connections (Fig 2b, *p* < 0.05). The classification accuracy of unconnected weights was high in the non-stimulated case, and therefore did not improve significantly with stimulation. The accuracy of predicted weights for all connection types improved with increasing the number of stimuli (Supplementary Fig. 1), particularly for strong inhibitory connections (Fig. 2c). The classification accuracy for inhibitory and excitatory connections improved drastically with increasing the number of stimuli, due to the activation of silent neurons (Fig. 2d)

### 3.3 Large neuronal populations

To determine the data requirements for deriving connection weights from larger neuronal populations, we calculated the average spike rate as an estimate of the number of spikes in the data from populations of sizes ranging from 200 to 2000 neurons (Fig. 4). The average spike rate increased linearly with the number of stimulations, and the slope of that relationship decreased with network size (Fig. 4a). The number of stimuli necessary to yield a firing rate similar to that in the 200 neuronal population with 15 stimuli increased linearly with population size (Fig. 4c). We next estimated the degree coverage afforded by the stimuli in populations of different size, as measured by the percentage of neurons that get stimulated by the random stimuli overall (Fig 4b). The number of stimuli necessary to get 99% coverage increased linearly with network size (Fig 4c), so that less than 200 stimulations were sufficient to get a full coverage for all the populations examined.

**Figure 4.**
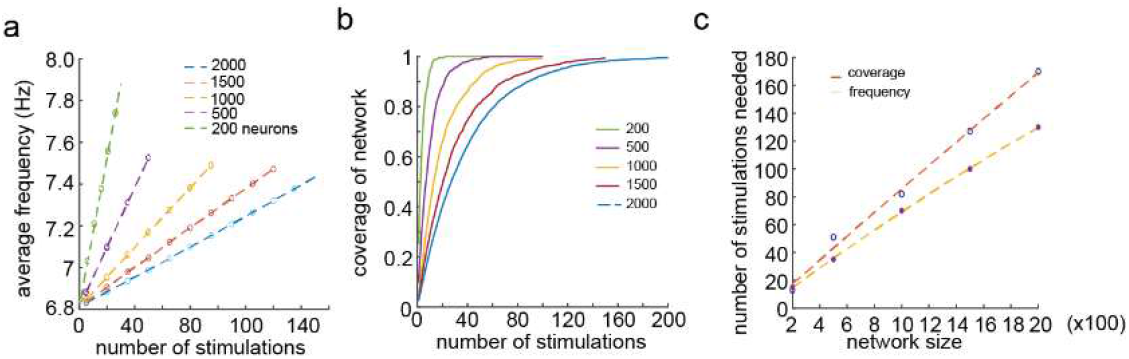
Deriving connectivity for large scale networks. **(a)** Average spike frequency as a function of the number of stimulations, for different network sizes. The frequency increased linearly with the number of stimulations. The slope decreased as the network size increased. **(b)** Coverage of the population by stimulation, measured by percent of the population activated, as a function of stimulus number. The slope of the curve decreased as the size of network increased. **(c)** The number of stimulations needed to reach 99% coverage increased linearly with network size. Similarly, the number of stimulations needed to reach a frequency of 7.4 Hz (corresponding to the 200 neurons, 15 stimuli case) increased linearly with network size.

## 4. Discussion

In this study, we demonstrated the utility of stimulation for deriving connectivity of neuronal populations with a significant proportion of the network comprising silent neurons, as seen physiologically. To date, the challenge imposed by silent neurons on deriving connections has remained largely unexplored. Here we showed that a supervised learning algorithm combined with a small number of stimulations enabled inferring connection weights with high fidelity and predicting spike trains with high accuracy. We have characterized the challenge imposed by silent neurons for different types of connections and for different population sizes. Our results make testable predictions about the number and protocol of the required stimulations, which is expected to enhance future efforts for deriving neuronal connectivity that underlies brain function.

Implementation of the method for diverse electrophysiological datasets relies on using experimental parameters for selective and continuous stimulation of interconnected neuronal subpopulations. Multiplexed single spike acquisition of connected networks during stimuli is routine both clinically and preclinically over multiple hours during different forms of stimuli [15, 22, 25]. Robust changes in firing rates are prevalent during both sensory and electrical stimuli, but fine tuning of firing patterns with only modest changes in firing rates is also possible. Recent work demonstrates modulation of subtle features of cortical activity including phase locking and inter-spike intervals of subpopulations, with only modest changes in firing rates [54]. Future expansion of our method will involve non poisson spike distribution during stimulation that emphasizes changes in spiking patterns over increase in rate, in order to identify scenarios that sufficiently add to the quantity of information needed for deriving weights and predicting spikes. Moreover, our method allowed for subtle prediction of single spikes evoked by single presynaptic cells (Fig. 4e-f). This offers a way to reconstruct networks containing neurons with highly selective receptive field that can also respond to abstract stimuli, a cell type seen physiologically in animals and humans in the form of place cells and abstract concept cells [55].

Our tests involve bringing a portion of the population to above firing baseline as a basis for deriving connectivity. Some stimulation experiments, particularly involving deep brain electrical stimulation for neurotherapeutic purposes, result in reduction of firing rates towards complete inhibition in some regions, and sometimes generates a combination of excitation and inhibition in response to stimulus [8, 56]. The paradigm we present here assumes targeting of a brain region excitatory to the network studied. This can in turn drive the design of new experiments for characterizing meso-scale connectivity of anatomical regions in the brain and while also facilitating maximization of the amount of information acquired about the network. Recent large scale recordings in awake rodents provide a viable platform for stimulus-derived connectivity mapping, demonstrating the activation of subpopulations across multiple brain regions, spanning the primary visual cortex, hippocampal and thalamic regions in response to constant visual stimulus [29]. Such increasingly accessible large-scale tools for electrophysiology will allow for methodically locating excitable regions at single-cell and single-spike resolution across the brain. Consequently, our results affirm that local electrical stimulation is not likely to facilitate network reconstruction, due in large part to indiscriminate synchronous firing and false positive observations that lead to imprecise determination of synaptic weights (supplementary Fig 3). This coincides with known constraints related to the symmetry of the radius of influence surrounding an electrode [57] that results in indiscriminate stimulation at the recorded region, and will likely obscure connectivity manifested in spiking patterns. For preclinical research, however, where optogenetic stimulation is available at the single cell type level [58], our results should lend themselves well where there is a high degree of control of the number and volumetric distribution of stimulated neurons. For reconstruction of large-scale networks, we find that longer stimulation sessions are required with a linear relationship between network size and number of stimulation epochs for sufficiently precise derivation of synaptic weights (Fig. 4). Long-term changes in network connectivity during training over many days [59] suggests that future studies involving very large scale electrophysiological recording and long stimulation sessions, network habituation and rewiring will have to be included in the derivation algorithm.

We characterized the use of stimulation to derive connections from a population of neurons onto a postsynaptic neuron. This enabled a focused investigation of the issue of silent neurons in deriving connections. Our results and stimulation framework can be used to expand the investigation into deriving connections from recurrent networks and overcome additional issues such as spurious connections due to spike correlations[60]. We employed an integrate and fire neuron model with simple implementation of synaptic connections. While such models are used widely to model brain networks[61], it will be of general interest to apply our stimulation framework to derive connections from groundtruth spiking data obtained from networks with detailed biophysical models of neurons and synaptic dynamics[50, 62, 63]. We anticipate that overcoming the non-linear challenges imposed by realistic neuronal networks will involve using our framework in tandem with methods for deriving connections based on spike cross-correlations[15, 22], or by using general purpose optimization methods such as genetic algorithms or maximum-likelihood[16, 64].

## 5. Conclusion

We established stimulation parameters for deriving connectivity of silent neuronal populations by way of a supervised learning algorithm. Our algorithm provides network reconstruction with high fidelity with accurate spike prediction. We have characterized performance for different connection types and population sizes, and established a method which is expected to enhance future efforts for deriving neuronal connectivity underlying brain function.

## Acknowledgements

This research was funded by NIH grant K01EB027184 and DP2NS122605 to AH. We thank Dr. Etay Hay for supplying reference code and useful comments on the manuscript.

## Author Contributions

XR performed the research. XR and AH wrote the manuscript.

## Competing interests

The authors declare no competing interests.

## Data availability

Datasets and associated figures (Figs. 1-4 and Supplementary Figs. 1-3) are available from the authors upon reasonable request.

## Notes

### Competing Interest Statement

The authors have declared no competing interest.

