## Supplemental Information for "Stimulation-mediated reverse engineering of silent neural networks"

### Supplementary information

#### Supplementary figures:

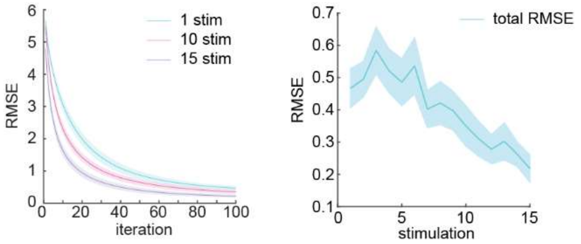


**Supplementary figure 1. Stimulation-dependent precision of derived weights**. (a) average RMSE of derived weights decreased rapidly within less than 100 iterations. (b) Larger number of stimuli resulted in more accurate derived weights.

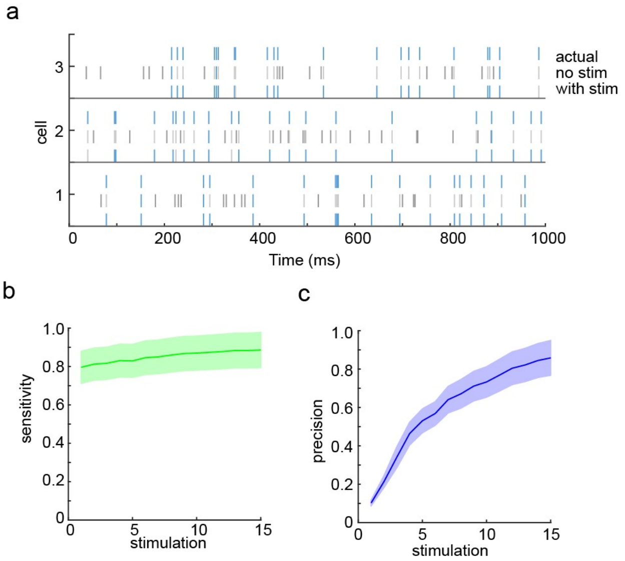


**Supplementary figure 2. Spike prediction using the derived connection weights for fully active network condition (a)** Three examples of spike trains of the postsynaptic neuron in the ground-truth test data (top), predicted spikes using derived weights in the non-stimulated population (middle), and stimulated population (bottom). Blue spikes are true positives. Light

[1] Department of Biomedical Engineering, University of Wisconsin-Madison. [2] Department of Electrical & Computer Engineering, University of Wisconsin-Madison.

grey spikes are false positives. Dark grey spikes are false negatives **(b)** Sensitivity of spike prediction as a function of stimulus number. The shaded area around the curve shows the standard deviation (n = 10). **(c)** Precision of spike prediction as a function of stimulus number.

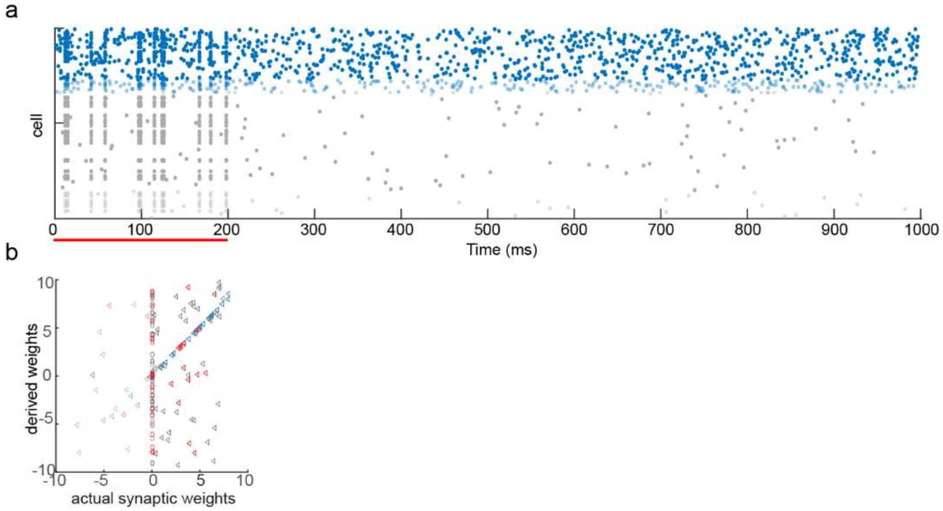


**Supplementary figure 3. Deriving synaptic weights using indiscriminate synchronized stimulation**. **(a)** Spike raster of the population spike trains. The top 34% of the trains correspond to active neurons and the bottom 66% correspond to silent neurons (see Methods). Stimulation was applied synchronously and is shown by the red bar. **(b)**. Derived weights from a population stimulated synchronously did not match the actual weights (r = 0.34, p = ~0).
